# Cholinergic basal forebrain neurons regulate vascular dynamics and cerebrospinal fluid flux

**DOI:** 10.1101/2024.08.25.609536

**Authors:** Kai-Hsiang Chuang, Xiaoqing Alice Zhou, Ying Xia, Zengmin Li, Lei Qian, Eamonn Eeles, Grace Ngiam, Jurgen Fripp, Elizabeth J. Coulson

## Abstract

Waste from the brain is cleared via a cerebrospinal fluid (CSF) exchange pathway, the dysfunction of which is suggested to underlie the pathogenesis of many brain conditions. Coherent cerebrovascular oscillation that couples with pulsatile CSF inflow is suggested to drive the fluid flux. However, how this coupling is regulated, whether it mediates waste clearance, and why fluid flux is impaired in disease status remain unclear. Here we show that vascular-CSF coupling correlates with cortical cholinergic activity in non-demented aged humans. The causal role of basal forebrain cholinergic neurons that project to the cortex is then verified by specific lesioning in mice, revealing correlated changes in vascular-CSF coupling, arterial pulsation and glymphatic flux, which can be altered by an acetylcholinesterase inhibitor. These results suggest a neurovascular mechanism by which CSF/glymphatic flux is modulated by cholinergic neuronal activity, thereby providing a conceptual basis for the development of diagnostics and treatments for glymphatic dysfunction.

## Introduction

Deficiency in the clearance of waste from the brain has recently been suggested to contribute to cognitive decline in aging, as well as the pathogenesis of a plethora of neurodegenerative diseases ^1^. Several routes, including the meningeal lymphatics, intramural periarterial drainage and the glymphatic system, have been hypothesized to support this clearance ^2–4^. The glymphatic system is an integrated part of the neurovascular unit by which waste in the interstitial fluid (ISF) can be removed from the brain. Driven by arterial pulsation, coherent neural and vascular oscillations, and other pressure changes ^5–8^, cerebrospinal fluid (CSF) moves along the periarterial space and influxes into the brain parenchyma, a process that is partly mediated by the aquaporin-4 (AQP4) water channels in the perivascular astrocytic end-feet. It then interchanges with solutes and macromolecules in the ISF, before passing into the perivenous space ^4^. Both human and animal studies have shown that impaired CSF-ISF drainage in aging ^9^ or sleep disorders ^10^ result not only in cognitive impairment ^1^^1^ but also in the accumulation of toxic molecules such as amyloid-b (Ab) ^4^, hyperphosphorylated tau ^1^^2,13^ and a-synuclein ^14^ aggregates, which are characteristic of neurodegenerative diseases. Reduced glymphatic flow has therefore been suggested to contribute to the pathogenesis of Alzheimer’s disease (AD), Parkinson’s disease and other neurological diseases ^1,15^.

Most work to date has focused on the mechanisms by which CSF and solutes move between the perivascular space and the brain parenchyma, or the consequences of dysfunctional glymphatic clearance. Several physiological factors can drive or affect glymphatic flow, including the perivascular pressure generated by arterial pulsation ^5,6^ and vasomotion ^16^, heart rate ^17^, hypertension ^6^ and state of arousal ^18^. Neural activity can also drive regional glymphatic flux via neurovascular coupling, as inhibition of vascular smooth muscle or neuronal activity abolishes the flux ^8,19^. Studies on anesthesia and sleep suggest that a strong glymphatic flow depends on coherent electrophysiological oscillations, in particular delta activity ^17,18^. This brain-wide coactivity can drive concorded vascular contraction, measured by global blood oxygenation level-dependent (BOLD) functional magnetic resonance imaging (fMRI) signal, leading to strongly anticorrelated CSF inflow in the fourth ventricle (BOLD-CSF coupling) ^7^. However, how ventricular CSF inflow relates to tissue glymphatic flux remains unclear. Due to the impact of sleep on glymphatic activity, it has been suggested that sleep disturbance contributes to glymphatic impairment in AD ^1^. Alternatively, disrupted CSF production, AQP4 function, vascular integrity and size of the perivascular parenchymal border macrophages, and periarterial pial layer may also lead to glymphatic deficits in aging and AD ^13,20–22^. However, the mechanism by which glymphatic flow is normally regulated and why it is impaired in AD are still poorly understood.

Here, we propose a neural mechanism of glymphatic dysregulation. The early and progressive loss of basal forebrain cholinergic neurons (BFCNs) is an important feature of AD that contributes to cognitive decline ^23^. BFCNs also play an important role in cerebrovascular regulation and neurovascular coupling by innervating cerebral arteries, capillaries, veins and interneurons to regulate vascular tone ^24^ and regional cerebral blood flow (CBF) ^25^. Intriguingly, BFCN loss and basal forebrain atrophy are associated with Ab burden in humans ^26,27^ and can exacerbate Ab pathology in animal models of dementia ^28–30^. However, the mechanism by which this occurs remains unclear. Recent studies have indicated that oscillation in the resting-state cortical BOLD signal correlates with basal forebrain activity, whereas inhibiting basal forebrain activity reduces BOLD oscillations ^31,32^. Given this neurovascular function, we questioned whether degeneration of BFCNs impairs glymphatic flux. We hypothesized that BFCN dysfunction would disrupt the BOLD-CSF coupling in regions receiving cholinergic projections, thereby reducing regional glymphatic flux.

To test this hypothesis, we conducted positron emission tomography (PET) and MRI scans in aged human subjects to assess the relationship between BOLD-CSF coupling and the cortical cholinergic density measured by a PET radiotracer, ^18^F-fluoroethoxybenzovesamicol (^18^F-FEOBV), which binds to the vesicular acetylcholine transporter and represents the presynaptic cholinergic nerve terminal density ^33^. We also verified the causal effects of cholinergic deficits on vascular and glymphatic dynamics in mice by targeted cholinergic lesion and treatment with an acetylcholinesterase inhibitor (AChEI). Our results demonstrate that BFCNs modulate arterial pulsation and BOLD-CSF coupling, leading to a change in tissue glymphatic flux in regions receiving BFCN innervation. These findings suggest a neurovascular mechanism of glymphatic regulation via a neuromodulatory system, the degeneration of which impairs glymphatic function in aging and disease.

## Results

### BOLD-CSF coupling correlates with cortical cholinergic density in aged human subjects

To understand the influence of cholinergic dysfunction on the BOLD-CSF coupling in non-demented aged subjects, we recruited 25 volunteers (age: 60-90 years; female: n=13). Among those, 10 subjects were diagnosed as exhibiting early mild cognitive impairment (MCI). Concurrent ^18^F-FEOBV PET and fMRI scans were collected to measure cortical cholinergic synaptic density, CSF ventricular inflow and cortical BOLD signals respectively (Fig. 1a). The coupling between the CSF and BOLD signals was assessed using cross-correlation analysis (Fig. 1b). Anti-correlation was observed with a median lag time of −1 scan (corresponding to a repetition time [TR] of 2.68 s), indicating that BOLD oscillations preceded CSF pulsation, consistent with previous studies ^7,34,35^. Here, we used the correlation coefficient of this lag time as a measure of the BOLD-CSF coupling. We also calculated the optimal lag time that maximizes the anticorrelation between BOLD and CSF oscillations. This coupling strength correlated with the optimal lag time that maximizes the anticorrelation (r = 0.48, p = 0.013; Fig. 1b), suggesting that weaker coupling may be due to CSF inflow not closely following the vascular dynamics.

**Fig. 1.**
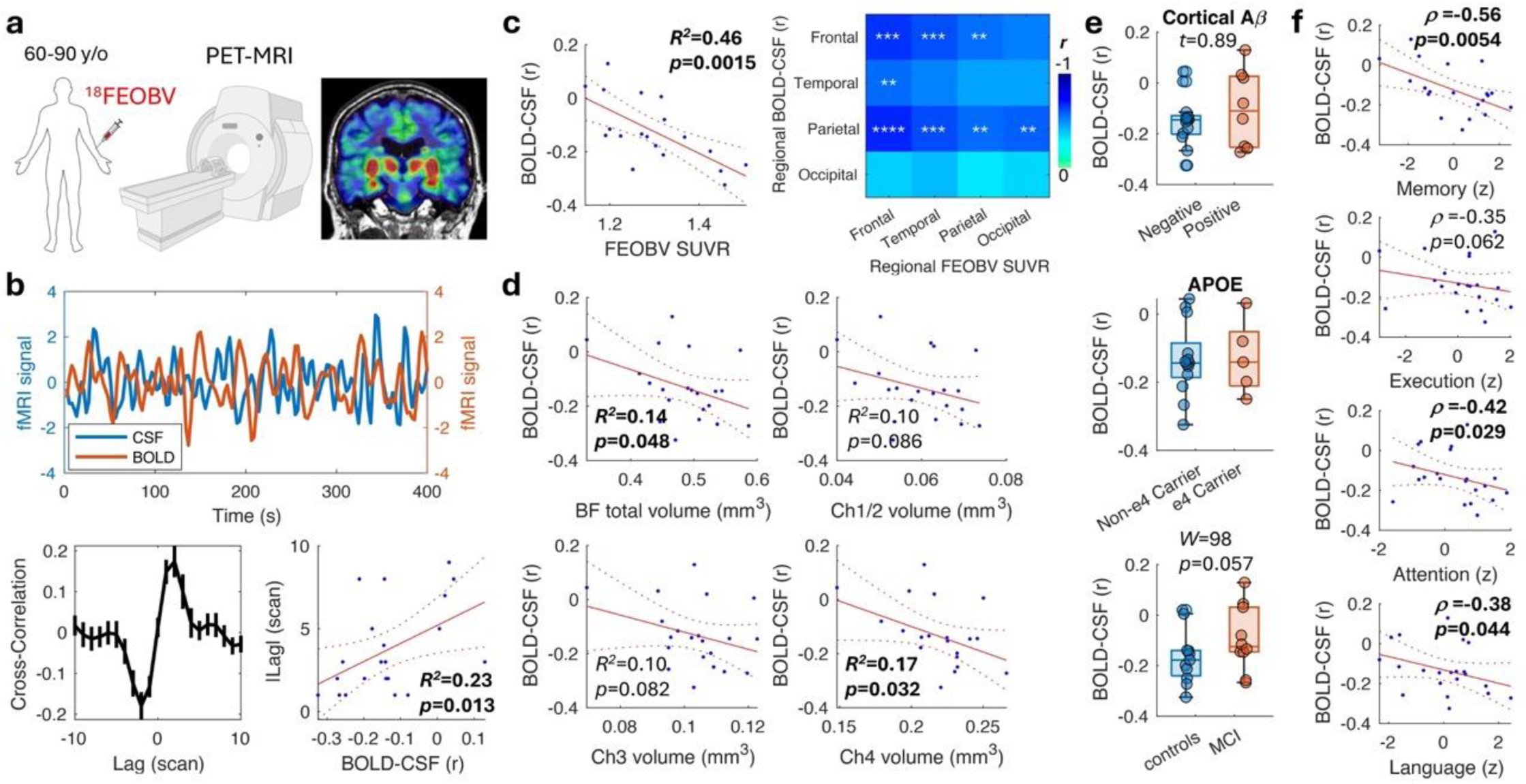
BOLD-CSF coupling correlates with cholinergic density in human. **a)** Concurrent PET-MRI with ^18^FEOBV tracer was used to measure cortical cholinergic vesicle density and resting-state BOLD signal in non-demented elderly subjects. The color-coded map shows an ^18^FEOBV PET image overlaid on structural MRI. **b)** An example of cortical BOLD and CSF inflow signals from a representative subject. The averaged cross-correlation between BOLD and CSF signals shows anticorrelation at a median time lag of −1 scan (TR = 2.68s per scan). The error bar represents the SEM. The BOLD-CSF coupling strength was defined as the Pearson correlation at lag = −1. This strength correlated with the lag time of the strongest anticorrelation in each subject. **c)** BOLD-CSF coupling correlated with total? cortical FEOBV SUVR. The matrix shows the correlation between regional BOLD-CSF coupling and regional cortical FEOBV SUVR. **: p<0.01; ***: p<0.005; ****: p<0.001 uncorrected. **d)** The BOLD-CSF coupling correlated with basal forebrain (BF) total and Ch4 but not other BF subregional volumes. **e)** No difference in the BOLD-CSF coupling was found between subjects of different cortical Aβ burden (top), APOE4 status (middle) and cognitive impairment (bottom). *W*: Wisconsin rank-sum test. **f)** The BOLD-CSF coupling of participants correlated with their overall neuropsychological scores in memory, attention and language. ρ: Spearman correlation coefficient. In the scatter plot, the red solid line represents the linear regression and the red dot lines represent the 95% confidence interval.

We found the BOLD-CSF coupling significantly correlated (r = −0.68, p = 0.0015, one tail) with the cortical cholinergic vesical density as measured by the FEOBV standardized uptake value ratio (SUVR) in the cortex, defined using structural MRI (Fig. 1c). This indicates that the lower the cholinergic activity in the cortex, the more disrupted the BOLD-CSF coupling became. The BOLD-CSF coupling was comparable between genders and across age or year of education, but was moderately correlated (r = 0.41, p = 0.033) with the volumes of white matter hyperintensities (Extended Data Fig. 1a). This suggests that the presence of small vessel disease may affect the coupling, consistent with a previous report ^36^. When the volume of white matter hyperintensity was controlled using partial correlation, the BOLD-CSF coupling still correlated with the cortical cholinergic density (r = −0.60, p = 0.0044). We further segmented the cortex into the frontal, temporal, parietal and occipital lobes to evaluate whether the relationship persisted in each region. The BOLD signal in each region was used to calculate the regional BOLD-CSF coupling. Our results revealed significant correlations between regional FEOBV SUVR and regional BOLD-CSF coupling in the frontal and parietal lobes, whereas the relationship was weaker in the temporal lobe and absent in the occipital lobe (Fig. 1c and Extended Data Fig. 1b), possibly due to the much weaker cholinergic FEOVB signal in the occipital lobe (F = 13.6, p = 2.9×10^-7^, one-way ANOVA; Extended Data Fig. 1c).

To verify that the changes in cholinergic density and BOLD-CSF coupling depend on BFCNs, we measured their relationships with the basal forebrain subregional volume from the structural MRI scans. As the nucleus basalis of Meynert (Ch4) of the basal forebrain is the primary source of cortical cholinergic innervation, we predicted that the Ch4 volume would correlate with cortical cholinergic density and BOLD-CSF coupling. Indeed, the Ch4 volume had the highest correlation with the cortical FEOBV SUVR (r = 0.60, p = 0.0020) in comparison with the other subregions, including Ch1/2 (medial septum [MS] and ventral diagonal band [VDB]) and Ch3 (horizontal diagonal band; Extended Data Fig. 1d). Moreover, the BOLD-CSF coupling significantly correlated only with the Ch4 volume (r = −0.41, p = 0.032; Fig. 1d). This indicates that the greater the cholinergic innervation from the basal forebrain, the stronger the coupling between cortical vascular dynamics and CSF inflow.

### BOLD-CSF coupling correlates with cognition but not amyloid burden

It has been reported that reduced BOLD-CSF coupling is associated with Aβ load in MCI and AD patients ^34,35,37^. We measured cortical Aβ burden (calculated using the Centiloid scale ^38,39^) using ^18^F-florbetaben PET with a threshold of 20 Centiloid to define abnormal levels of Aβ. The BOLD-CSF coupling was comparable between the Aβ-positive and -negative subjects (Fig. 1e), and there was no correlation with the Aβ burden (R^2^ = 0.04, p = 0.18; Extended Data Fig. 1e). The BOLD-CSF coupling was also not associated with apolipoprotein E (APOE) genotype. However, there was a trend of weaker BOLD-CSF coupling in subjects classified as MCI (W = 98, p = 0.057, Wisconsin rank-sum test; Fig. 1e). Although there was no correlation with the Mini-Mental State Examination (MMSE) score, the BOLD-CSF coupling correlated with three out of the four cognitive domains, memory, executive function, attention and language, in the neuropsychological assessments (Fig. 1f; see Extended Data Fig. 1f for comparisons with specific test). This cognitive correlation likely reflects a contribution by the cortical cholinergic vesicle density, as it became insignificant when the FEOBV level was controlled.

### Cholinergic lesion reduces BOLD-CSF coupling in mice

To test whether BOLD-CSF coupling is causally regulated by basal forebrain cholinergic innervation, we selectively ablated cholinergic neurons in the MS and VDB that project to the hippocampus ^40^ to restrict the resultant effects in this target area. As BFCNs are marked by the expression of the p75 neurotrophin receptor (hereafter referred to as p75), we injected a murine-p75 antibody conjugated to the immunotoxin saporin or an IgG-saporin as sham control into the MS/VDB of young healthy wild-type mice. This resulted in a significantly reduced p75^+^ neuron number in the MS and VDB (Fig. 2a) of p75-saporin-injected mice, confirming the presence of BFCN degeneration.

**Fig. 2.**
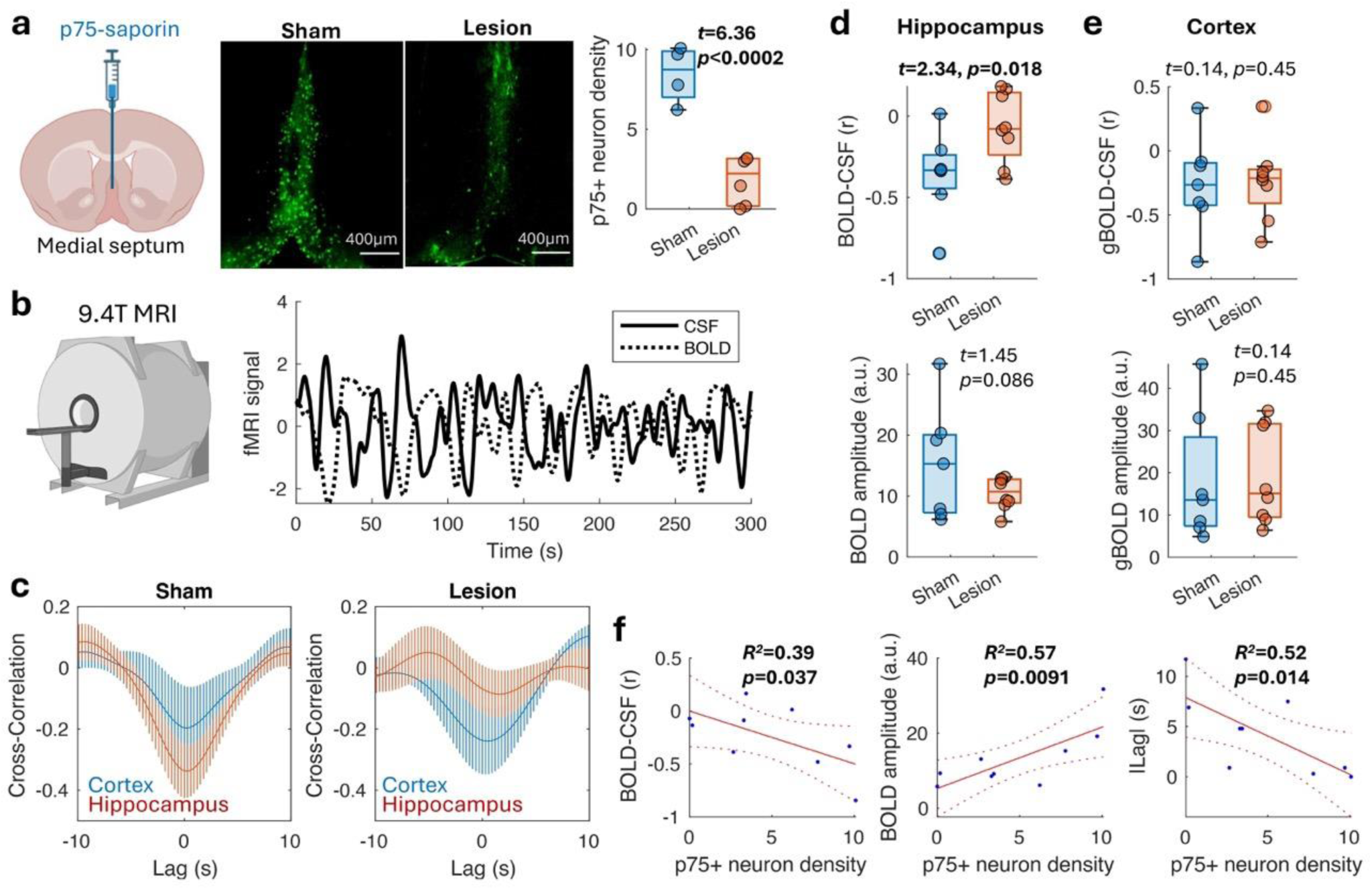
Septal cholinergic lesions reduce hippocampal BOLD-CSF coupling in mice. **a)** An immunotoxin, p75-saporin, which targets the p75 neurotrophin receptor-expressing cholinergic neurons was injected into the medial septum (MS) of the mouse basal forebrain. This resulted in significant reduction of cholinergic neurons (green fluorescence) and their density in the MS compared to that of sham mice injected with the non-targeted IgG-saporin. **b)** 3-4 weeks after the surgery, resting-state fMRI was conducted using a 9.4T MRI to measure the cortical BOLD (dash line) and ventricular CSF inflow (solid line) signals. **c)** The cross-correlation between the cortical BOLD and CSF signals shows anticorrelation at 0 lag time in both sham and lesioned mice. However, the anticorrelation between the regional BOLD and CSF signals were abolished in the lesioned mice in the hippocampus, which receives the MS cholinergic projection, but not in the cortex. Error bars represent the SEM. **d)** The cortical BOLD signal coupling with the CSF signal (top) and fluctuation amplitude (bottom) were comparable between the lesioned and sham mice. **e)** In the hippocampus, however, the hippocampal BOLD-CSF coupling strength was reduced toward 0, with the BOLD signal amplitude showed a trend of decrease. **f)** The hippocampal BOLD-CSF coupling, BOLD signal amplitude, and the lag time of maximal anticorrelation between the BOLD and CSF signals, were all significantly correlated with the p75-positive cholinergic neuron density of the MS of the imaged mice.

With the cholinergic inputs to the hippocampus denervated, we predicted that the hippocampal BOLD signal would be disrupted. To test this, we conducted resting-state fMRI in mice using an ultrafast imaging (temporal resolution = 0.3 s) to capture the fluid dynamics under a mixture of medetomidine sedation and light isoflurane anesthesia ^41^. Following similar data preprocessing to that used for human scans, the ventricular CSF inflow signal was measured from the aqueduct and compared with the BOLD signal in the cortex and hippocampus. We found a similar anticorrelation between the CSF and BOLD oscillations in sham control mice (n = 7) to that seen in humans (Fig. 2b). However, different from the findings in humans, virtually no lag time was found in either the cortex or hippocampus in the control mice (indicative of a more instant anticorrelation) using cross-correlation analysis (Fig. 2c). This finding may be due to the very fast heart rate and small brain of the mouse, which permit tighter physiological responses.

In the cholinergic lesioned mice (n = 8), the anticorrelation was greatly reduced, trending towards no association, in the hippocampus (t = 2.34, p = 0.018, two-sample t-test, one tail; Fig. 2d) whereas the cortical BOLD signal remained anticorrelated with the CSF signal (Fig. 2e). The optimal lag time for maximizing anticorrelation between BOLD and CSF signals also strongly correlated with hippocampal BOLD-CSF coupling (r = 0.77, p = 0.0004), confirming the observation in humans. Comparing the amplitude of BOLD oscillations, calculated based on the temporal standard deviation, the lesioned mice also displayed a trend of reduced amplitude in the hippocampus (t = 1.45, p = 0.086; Fig. 2d) but not in the cortex (Fig. 2e). This is consistent with a previous study which reported that inhibition of the basal forebrain reduces the amplitude of the BOLD signal ^31^.

Importantly, the hippocampal BOLD-CSF coupling correlated with the p75^+^ cholinergic neuron density in the MS (r = −0.62, p = 0.037; Fig. 2f), corroborating our FEOBV findings in humans. The BOLD oscillation amplitude and optimal lag time also correlated with the MS cholinergic neuron density (r = 0.75 and −0.72, respectively; Fig. 2f). This region-specific and neuronal density-correlated disruption of the BOLD signal and its coupling with CSF inflow indicate that cholinergic innervation is important for modulating targeted brain regional vascular dynamics and that cholinergic deficits can uncouple CSF inflow from the regional neural and vascular demands.

### Cholinergic lesion alters glymphatic flux in mice

Although the coupling of vascular and CSF oscillations has been suggested as a mechanism of glymphatic influx, whether this leads to a change in tissue glymphatic flux remains unknown. Two-photon microscopy has been used to delineate CSF transport along the pial artery and in tissue; however, it is difficult to use this technique to image deep structures such as the hippocampus without compromising the cortical tissue and fluid pressure. In order to measure the glymphatic flux over the whole brain, we therefore used small molecule gadolinium (Gd) contrast-enhanced MRI ^42^ in a subset of mice (lesion: n = 6, sham: n = 4). This approach was conducted with slow infusion to minimize the impact on intracranial pressure ^43^ under the same medetomidine sedative protocol that has been shown to promote glymphatic flow similar to that during sleep ^44^.

Following intracisternal infusion (Fig. 3a), we observed that the Gd tracer entered the brain along the ventral-caudal axis into various brain regions as indicated from the signal enhancement in the dynamic 3D T1-weighted MRI scans. The percentage signal change during the time-course of the MRI scan was determined. To estimate the accumulated glymphatic flow volume, we calculated the signal change area under the curve (AUC) for each voxel. After co-registering all the AUC maps to a brain template, voxel-wise comparison of the results from the lesioned mice with those of the sham control group revealed an increase in Gd contrast in limited brain regions but including the hippocampus (t = 2.01, p = 0.040; Fig. 3b). We also mapped the correlation coefficient between MS cholinergic neuron number and the AUC in each voxel. Our results revealed a negative relationship with the number of these MS cholinergic neurons in various regions, including the ventral hippocampus, amygdala, thalamus and basal forebrain (Fig. 3c). As an increased Gd AUC is typically interpreted as greater glymphatic flux, this finding seemed to contradict the disrupted BOLD-CSF coupling under cholinergic lesion.

**Fig. 3.**
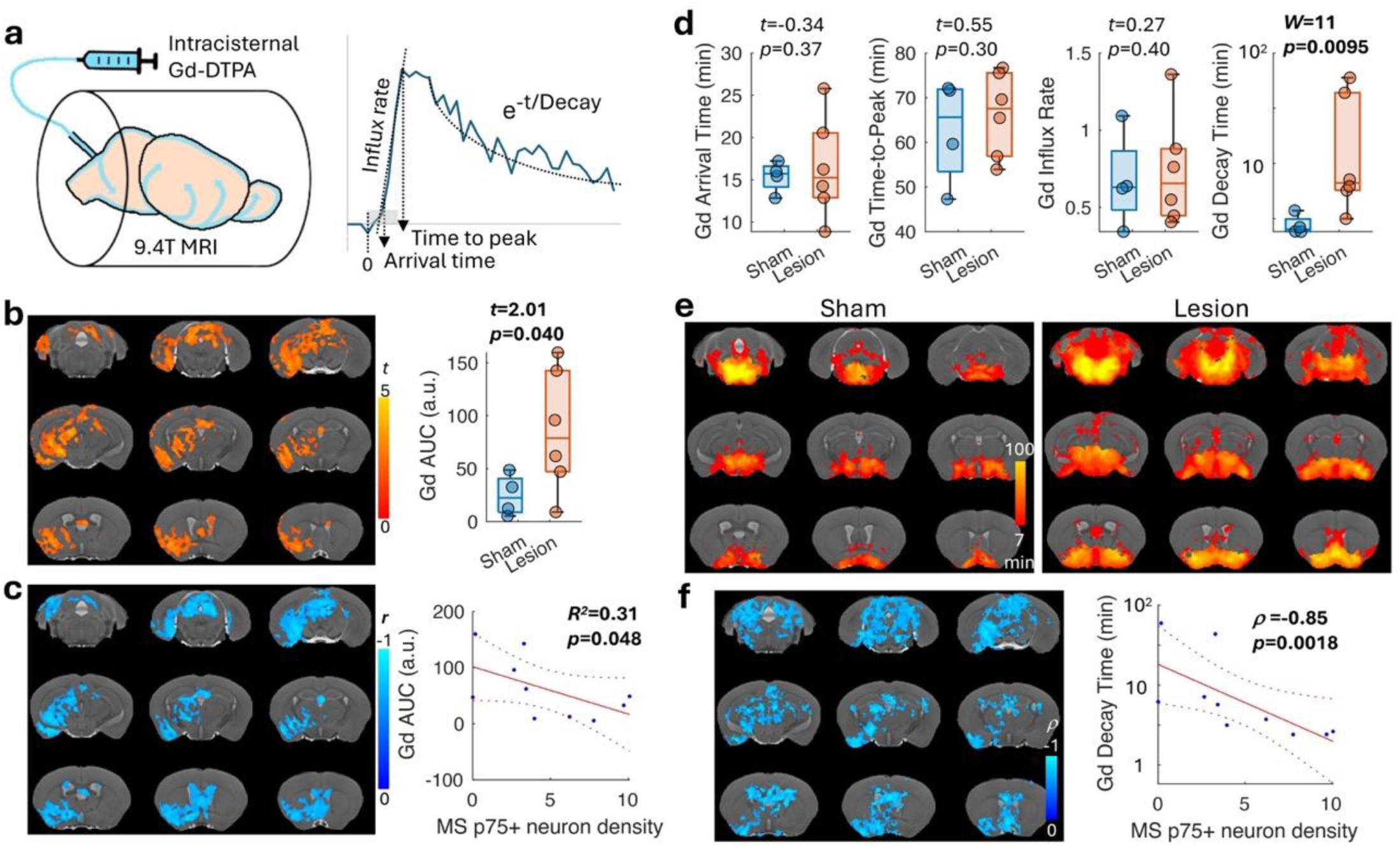
Septal cholinergic lesions affect hippocampal glymphatic flux. **a)** The glymphatic signal was detected by time-series T1-weighted 3D MRI scans with Gd-DTPA slowly infused into the cistern magna. Besides the area-under-the-curve (AUC, which represents accumulation), the Gd contrast arrival time, time-to-peak, rising slope (influx rate), and its exponential decay time (efflux) were calculated. The gray bar starting from t = 0 represents the duration of Gd contrast infusion. **b)** Two-sample t-test between the AUC maps of the lesion and sham mice shows significantly higher AUC in the lesioned mice (left; voxel-wise p<0.05, cluster-wise p<0.05 for correction of multiple comparison). When quantified (right), the AUC in the hippocampus was significantly higher in the lesioned mice. **c)** Significant negative correlation coefficient between the AUC and MS cholinergic neuron density can be found in similar brain regions as in b) (left; voxel p<0.05, cluster p<0.05). Furthermore, the AUC in the hippocampus significantly correlated with MS cholinergic neuron density (right). **d)** In the hippocampus, the Gd contrast arrival time, time-to-peak and influx rate were comparable between the sham and lesioned mice. However, the decay time was significantly longer in the lesioned mice. **e)** The group decay time maps (one-sample t-test, voxel p<0.05, cluster p<0.05) indicated overall slower (more yellow) efflux in the lesioned mice. **f)** The decay time negatively correlated with the MS cholinergic neuron density in the subcortical areas, particularly the midbrain (Spearman correlation, voxel p<0.05, cluster p<0.05). In the hippocampus, the decay time negatively correlated with the MS cholinergic neuron density. ρ: Spearman correlation coefficient.

To better characterize the kinetics of glymphatic flux, we next calculated the Gd contrast arrival time, time-to-peak, rising slope (as a measure of influx rate) and the signal exponential decay time (as an indication of efflux) in each voxel. Regional analyses showed that, among these kinetic measures, the decay time was significantly longer (W = 11, p = 0.0095, Wilcoxon rank-sum test, one tail) in the hippocampus of lesioned mice (Fig. 3d). Voxel-wise comparisons demonstrated that the Gd decay time was also generally longer in the lesioned mice (Fig. 3e). Further correlation with the MS cholinergic neuron count revealed a negative correlation in similar regions to those seen in the AUC mapping (Fig. 3f). Regional analysis also confirmed that cholinergic neuron density highly correlated with the hippocampal Gd decay time (ρ = −0.85, p=0.0018, Spearman’s correlation, one tail). This suggests that the larger Gd AUC in mice with cholinergic lesions was due to slower Gd efflux and hence resulted in more Gd retention in the hippocampal tissue. This anatomical and functional correspondence between glymphatic change and the known MS/VDB cholinergic projection to the hippocampus demonstrates a specific role of BFCNs in regulating glymphatic function in the innervated brain areas.

### BOLD-CSF coupling correlates with glymphatic flux

To determine whether coupling of the vascular oscillation and ventricular CSF inflow is related to the downstream tissue glymphatic flux, we compared the hippocampal BOLD-CSF coupling with the Gd contrast kinetics in the hippocampus. This revealed that the Gd contrast time-to-peak positively correlated with BOLD-CSF coupling (ρ = 0.87, p=0.0023; Fig. 4a), indicating that the tighter the BOLD-CSF coupling, the faster CSF/glymphatic influx occurs. We also found that a shorter lag time with ventricular CSF inflow (Fig. 4b) and a larger BOLD oscillation amplitude (Fig. 4c) lead to faster glymphatic influx. This suggests that regional BOLD-CSF coupling mediates tissue glymphatic influx. Interestingly, the lag time and BOLD amplitude also showed trends of correlation with the Gd decay time (Fig. 4b and c), implicating a potential effect on the glymphatic efflux as well.

**Fig. 4.**
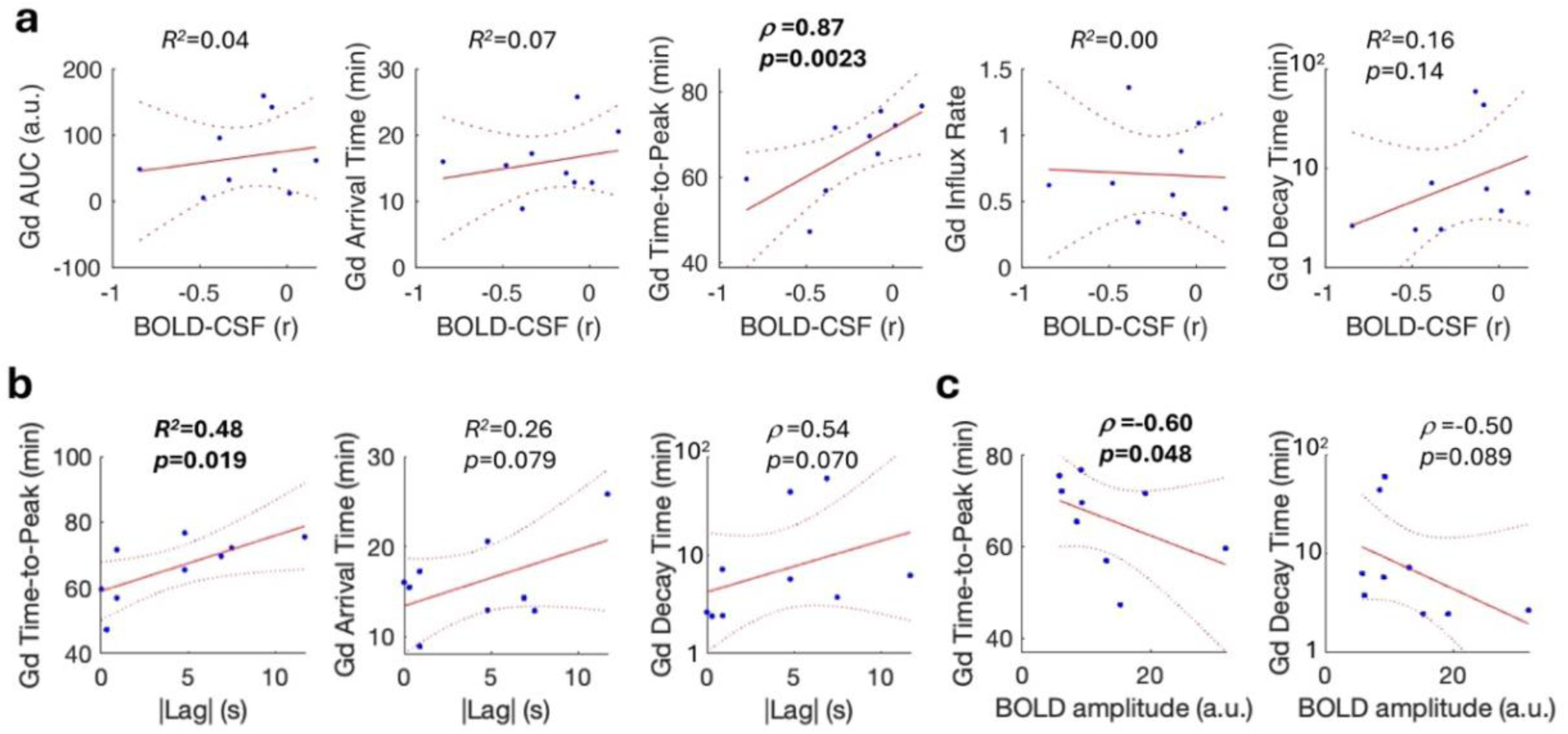
BOLD-CSF coupling correlates with glymphatic flux in the mouse hippocampus. **a)** Comparing the hippocampal BOLD-CSF coupling with the hippocampal glymphatic signal measured by intracisternal Gd contrast shows significant correlation with the time-to-peak and a trend of correlation with the decay time. **b)** The lag time for maximal anticorrelation between hippocampal BOLD and CSF signals positively correlated with the time-to-peak and marginally correlated with the arrival time and decay time. **c)** The hippocampal BOLD signal fluctuation amplitude negatively correlated with time-to-peak and marginally correlated with the decay time. ρ: Spearman correlation coefficient.

### Cholinergic lesion reduces arterial pulsation

As BFCNs are involved in the modulation of cortical activity ^45^, vascular tone ^24^ and the sleep-awake cycle ^46^, these factors could co-contribute to the observed glymphatic change. Based on a previous study which demonstrated that cholinergic ablation denervates the arteries and arterioles ^47^, we hypothesized that MS/VDB cholinergic neurons regulate the arteries that feed into the hippocampus. To test this, we used magnetic resonance angiography to distinguish the major cerebral arteries in each mouse (Fig. 5a), particularly the posterior cerebral artery (PCA) and the proximal longitudinal hippocampal artery (LHiA) that supply the hippocampus ^48^. To measure arterial pulsation in deep brain structures, we used the flow-related enhancement of gradient-echo MRI, in which the signal change is proportional to the flow velocity when the imaging is faster than the pulsatile flow across the imaging slice ^49^ and the same effect used for detecting CSF inflow. As the arterial blood flow velocity is proportional to the pressure wave velocity, which in turn is proportional to the square root of the vessel diameter ^50^, the arterial flow velocity represents a surrogate measure of the arterial wall movement (arterial pulsation). In this imaging, faster flow velocity leads to a higher signal ^49^ that follows the arterial pulse pressure changes ^51^. This was verified by comparing the MRI signal changes with pulsation measured at the tail artery using a pulse oximeter (Fig. 5a). The pulsatile signals obtained at the PCA and LHiA exhibited a high spectral power at the cardiac pulse frequencies (including the aliased harmonics) and lower spectral power corresponding to the respiration rate and vasomotion (0.01-0.1Hz ^16^) range (Fig. 5a). We therefore filtered the MRI signal based on the primary cardiac frequency to calculate the arterial pulsation power.

**Fig. 5.**
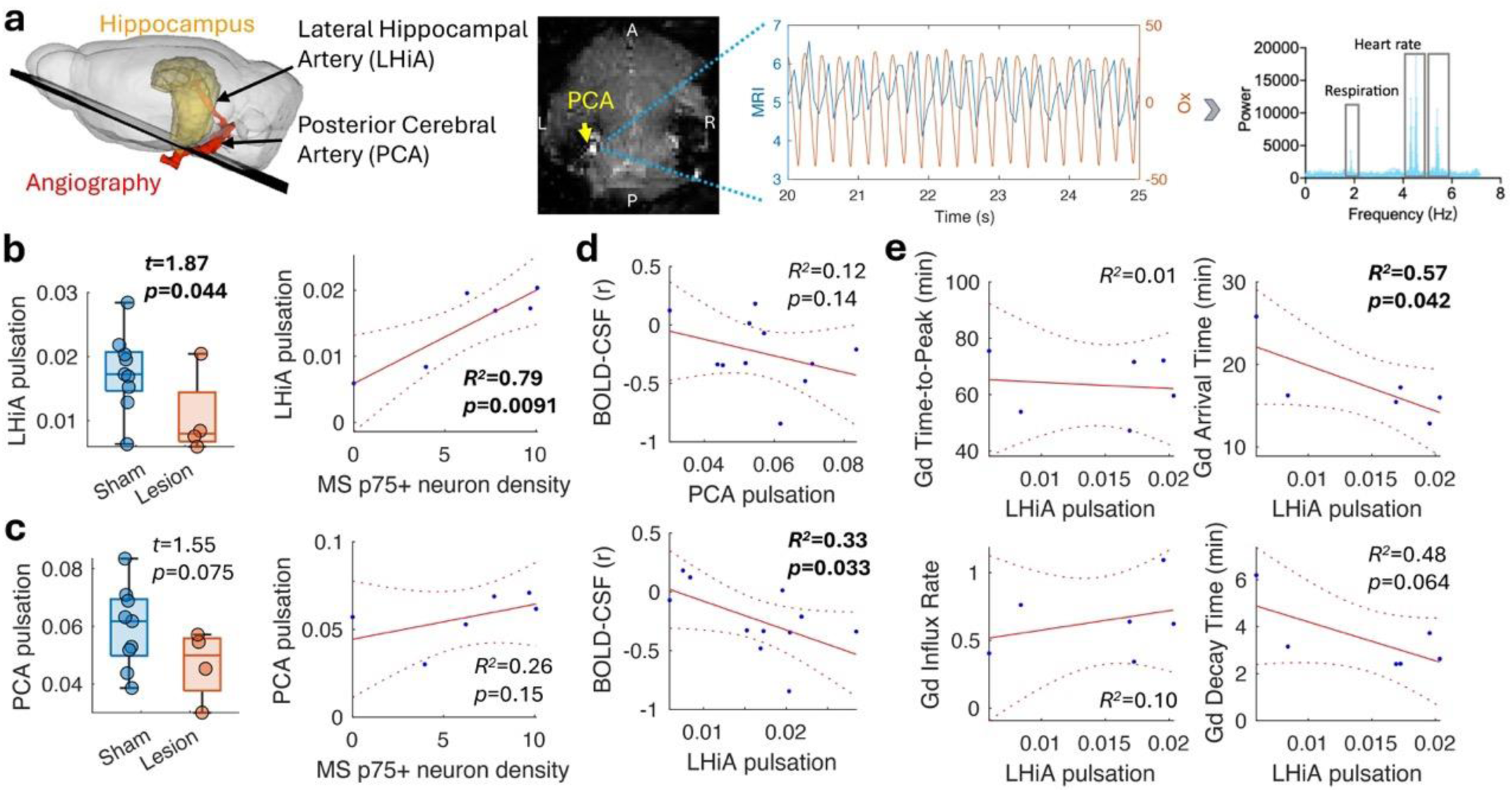
Cholinergic lesion attenuates arterial pulsation. **a)** Ultrafast MRI scans were used to detect the signal changes induced by the pulsatile blood inflow of large cerebral arteries. Compared with the pulsation of the tail artery measured by the pulse oximeter (Ox; red line), the MRI signal (blue line) closely follows the cardiac pulses (middle). Fourier transform of the MRI signal shows high peaks at the heart rate and lower peak at the respiration rate (right). **b)** The variance of MRI signal change around the cardiac frequency was used to estimate the arterial pulsation. Compared to the sham mice, the pulsation at the LHiA was significantly reduced in the lesioned mice and the pulsation correlated with MS cholinergic neuron density. **c)** Pulsation of the PCA was not sufficiently affected and was not significantly correlated with cholinergic neuron density. **d)** The hippocampal BOLD-CSF coupling correlated with LHiA pulsation but not with PCA pulsation. **e)** The LHiA pulsation significantly correlated with the Gd contrast arrival time and shows a trend of correlation with the decay time in the hippocampus.

In comparison with the sham controls (n = 9), mice with cholinergic lesions (n = 4) displayed greatly reduced arterial pulsation of the LHiA (t = 1.87, p = 0.044; Fig. 5b). The altered LHiA pulsation also strongly correlated with MS cholinergic neuron density (r = 0.89, p = 0.0091). In contrast, there was a trend towards a decrease for the PCA pulsation (t = 1.55, p = 0.075) which did not correlate with cholinergic neuron density (r = 0.51, p = 0.15; Fig. 5c). Stronger LHiA pulsation correlated with tighter hippocampal BOLD-CSF coupling (r = −0.57, p = 0.033; Fig. 5d) but not with BOLD oscillation amplitude (r = 0.36, p = 0.13), suggesting that upstream arterial pulsatility may not affect the tissue neurovascular response. Importantly, stronger LHiA pulsation correlated with shorter Gd contrast arrival time (r = −0.76, p = 0.042; Fig. 5e), which is consistent with the role of local arterial pulsation in mediating perivascular fluid transport. Moreover, mice with stronger LHiA pulsation tended to have a shorter Gd decay time (faster efflux; r = −0.69, p = 0.064; Fig. 5e), suggesting a potential effect of pulsatility on the efflux. In contrast, PCA pulsation did not correlate with regional BOLD-CSF coupling or glymphatic influx.

### BOLD-CSF coupling and arterial pulsation jointly mediate fluid flux

As pulsation of the feeding artery and downstream hemodynamics in tissues could have differential contributions to fluid transport, influx and efflux, we compared linear models using BOLD-CSF coupling and LHiA pulsation individually or jointly to predict glymphatic signal kinetics (Extended Data Table 1). Our results revealed that LHiA pulsation had a major contribution to the arrival time, whereas BOLD-CSF coupling had stronger effect on the time-to-peak, which reflects how fast most of the tracer is distributed within the tissue. The AUC and influx rate were jointly contributed by both pulsation and coupling strength. Interestingly, pulsation and coupling strength had rather independent contribution to the decay time. These results indicate that the observed altered glymphatic flux may be due to regional uncoupling from the CSF inflow and weakened perivascular transport caused by the cholinergic deficit in the lesioned mice. Considering that BFCN lesions do not alter sleep-awake cycle ^52^ or overall neural activity ^53^, our findings suggest a mechanism by which BFCNs regulate glymphatic flux in their projection areas through modulation of both the feeding arterial pulsation and tissue neurovascular synchronization (regional BOLD).

### Cholinergic treatment alters BOLD-CSF coupling, arterial pulsation and glymphatic flux

To test whether treatments that target the cholinergic system can normalize the dysregulated vascular dynamics and ISF/CSF flux, we used an AChEI, donepezil, which has been widely adopted to treat cognitive impairment in AD patients by compensating for reduced cholinergic neurotransmission due to neurodegeneration of BFCNs. We acutely delivered donepezil (2mg/kg, intraperitoneal [i.p.]; n = 6) or vehicle (saline; n = 6) into cholinergic lesioned mice (Fig. 6a), after which we conducted resting-state fMRI and arterial pulsation MRI, followed by cisternal injection of Gd contrast with dynamic T1-weighted MRI to measure glymphatic flux. Our results revealed a significant change in BOLD-CSF coupling in mice under donepezil treatment (t = 2.65, p = 0.012; Fig. 6b). Surprisingly, instead of normalizing towards stronger anticorrelation, the BOLD and CSF signals became positively correlated. Donepezil treatment also significantly reduced the PCA pulsation (t = −3.06, p = 0.0060) and marginally reduced LHiA pulsation (t = −1.13, p = 0.14; Fig. 6c). These vascular changes may be due to reduced pulsatility of fully dilated arteries ^54^ and cardiovascular effects of donepezil, as reflected in the altered heart rate we observed (t = 2.12, p = 0.030; Fig. 6c). Voxel-wise comparison of the Gd contrast AUC and kinetic parameters showed that donepezil treatment shortened the decay time in the ventral-caudal part of the brain, where labelled fluid is first seen to influx into the perivascular space around the major cerebral arteries, whereas the decay time in the anterior brain around the striatum (which contains inhibitory cholinergic interneurons) was prolonged (Fig. 6d). Although regional analyses revealed no change in the hippocampal glymphatic flux between treated and untreated lesioned mice (Fig. 6d), the influx rate again positively correlated with the LHiA pulsation (ρ = 0.58, p = 0.044; Fig. 6f) and marginally correlated with the BOLD-CSF coupling (ρ = −0.55, p = 0.052; Fig. 6e). Interestingly, the weaker LHiA pulsation in donepezil-treated mice correlated with faster hippocampal efflux measured by the Gd decay time (ρ = 0.56, p = 0.048), suggesting that the upstream vascular effects rather than within tissue neurovascular effects may have been the dominant response of the treatment.

**Fig. 6.**
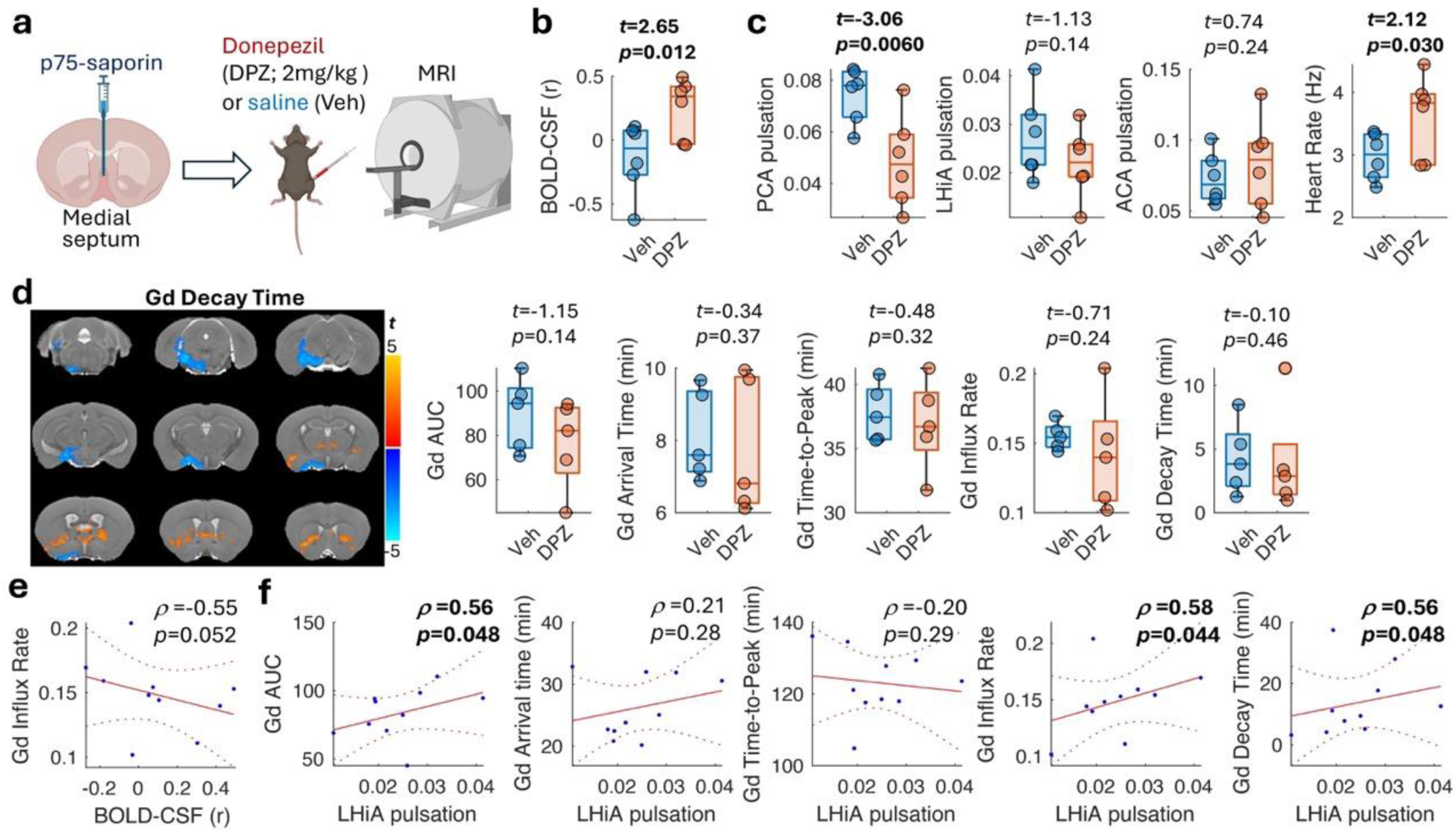
Donepezil treatment alters BOLD-CSF coupling and arterial pulsation but not glymphatic flux. **a)** Donepezil (DPZ) or the vehicle saline (Veh) was injected peritoneally in MS cholinergic lesioned mice. **b)** The hippocampal BOLD-CSF coupling was significantly increased and displayed a positive correlation in the DPZ-treated mice. **c)** DPZ treatment increased the heart rate, but significantly decreased the PCA pulsation and marginally reduced the LHiA pulsation, while the pulsation of the anterior cerebral artery (ACA) was unchanged. **d)** The glymphatic signal measured by Gd contrast shows that DPZ treatment led to a shorter decay time at the ventral part of the brain but longer decay time near the striatum (two-sample t-test, voxel p<0.05, cluster p<0.05). In the hippocampus, DPZ treatment did not change the glymphatic signal AUC or kinetics. **e)** However, the hippocampal BOLD-CSF coupling shows a trend of correlation with the Gd contrast influx rate. **f)** The pulsation of the hippocampal artery (LHiA) also correlated with the glymphatic signal AUC, influx rate and decay time.

Together these results suggest that the neurovascular regulation of BFCNs is an important factor for glymphatic function with cholinergic neurodegeneration impairs glymphatic flux. While AChEI treatment supports this mechanism, the intervention is not an effective means for improving glymphatic flux, consistent with donepezil not being a disease-modifying treatment for AD.

## Discussion

The regulation of glymphatic flow has been a subject of interest since the discovery that it has the ability to clear pathological protein aggregates from the brain ^4,13^ and is impaired in AD ^55,56^. Our findings demonstrate that BFCNs maintain normal CSF-ISF flux via neurovascular regulation. Using a cholinergic PET tracer, we identified an important role of BFCNs in coordinating cerebrovascular dynamics and CSF inflow in aged humans. The causal role of BFCNs and its relevance to glymphatic flux were further demonstrated in a mouse model of cholinergic neurodegeneration, revealing a strong correlation between BOLD-CSF uncoupling, reduced arterial pulsation, and glymphatic flux in BFCN-denervated brain areas, particularly the hippocampus. BFCN degeneration is an early feature of AD that correlates with and predicts cognitive impairment and Aβ burden ^27,45^. Our findings suggest a causative relationship between BFCN dysfunction and glymphatic deficiencies during aging and AD development.

We provide direct evidence that BFCNs regulate the BOLD-CSF coupling in awake human subjects and lightly anesthetized mice. The electrophysiological delta activity has been suggested as a neural mechanism of elevated BOLD-CSF coupling during sleep ^7^. Systemic factors, such as peripheral hemodynamic fluctuations ^57^, cerebrovascular response to deep breath ^58^ and cardiac-dependent vascular and CSF oscillations ^37^, have also been reported to contribute to BOLD-CSF coupling. BFCNs modulate both neural synchrony and neurovascular coupling, with lesion of cholinergic neurons being reported to largely attenuate local field potential and hemodynamic response to stimuli ^29,59,60^. This neurovascular effect could underpin the reduced BOLD signal and therefore its coupling with CSF inflow. The anatomically and functionally specific changes in this measure as a result of decreased cholinergic innervation support a neural origin for the coupling. We further confirmed that BOLD-CSF coupling directly relates to tissue glymphatic signal kinetics. This demonstrates that the macroscopic MRI measure could reflect tissue fluid flux, supporting the use of BOLD-CSF coupling as a non-invasive marker of glymphatic function.

We identified arterial pulsation as another key factor that contributes to glymphatic regulation by cholinergic neurons. Numerous studies have shown the impact of cholinergic neurodegeneration and AChEI treatment on CBF ^61–64^. Despite the effect on the tissue capillary blood flow, whether BFCNs also regulate the pulsatility of the artery being innervated is unknown. Our findings indicate that cholinergic activity modulates pulsation of large cerebral arteries, which correlates with the downstream tissue glymphatic signal. Previous microscopy studies reported that pulsation of penetrating arterioles mediates perivascular CSF inflow ^5,6^. Recent MRI studies in humans also demonstrated perivascular fluid change following cardiac pulses ^65,66^. Our findings extend these observations and link the pulsatility of upstream feeding artery with tissue glymphatic signal kinetics related to influx and efflux. As the venous vessels also present pulsatile variations but with a certain delay time from the artery ^67,68^, the attenuated arterial pulsation under BFCN denervation may reduce venous pulsation thus impacting the fluid efflux. Further study will be needed to elucidate the relationship with perivenous fluid flux.

AChEIs are widely prescribed to patients suffering cognitive impairment, and have been shown to alter neural oscillation and CBF in diverse brain regions in MCI and AD patients ^61–64^. We explored the use of an AChEI, donepezil, in modulating BOLD-CSF coupling, arterial pulsation and glymphatic flux. The minimal effects we observed on glymphatic flux may be due to its disruption of BOLD-CSF coupling and attenuation of arterial pulsation. Furthermore, donepezil not only reduces acetylcholine degradation throughout the body but also increases or decreases other neuromodulators, such as dopamine and norepinephrine, in different brain regions ^69^. The regionally dependent neural and vascular effects and the involvement of multiple neuromodulators complicates the possible effects that donepezil has on glymphatic function. Despite the lack of desirable efficacy, we found that pulsation of the hippocampal artery correlated with glymphatic flux after treatment. Future studies may titrate the dosage to minimize peripheral effects or focus on targeted modulation of arterial pulsation in order to enhance glymphatic function.

The glymphatic pathway is reliant on perivascular transport and trans-astrocytic bulk flow. BFCNs directly innervate the astrocytes (including the perivascular astrocyte endfeet ^47^), smooth muscle cells and endothelial cells of the neurovascular unit ^25^. Cholinergic innervation of this unit is known to be important for the maintenance of vascular tone and in the mediation of site-directed blood flow. Affecting the neurovascular unit by the BFCN lesioning method used herein ^47^ resulted in reduced arterial pulsation and its coupling with CSF inflow. Although we cannot rule out that inflammation and a disrupted blood-brain barrier induced by the cholinergic denervation may also contribute to the glymphatic dysfunction, the sham control mice would experience similar levels of dysfunction.

The current cholinergic-vascular hypothesis suggests that hypoperfusion and dysregulated neurovascular coupling due to BFCN degeneration contributes to cognitive decline and neurodegeneration in AD ^70^. Recent studies have reported that abnormalities in measures of perivascular flux correlate with neuropsychological performance and cerebral grey matter volumes in people with or at risk of AD ^56,71^. With basal forebrain atrophy shown to precede cortical degeneration and memory impairment in AD ^72^, this early BFCN degeneration could lead to glymphatic deficits that promote the accumulation of pathological molecules. Indeed, AD mouse models show accelerated Aβ pathology and disease progression following BFCN lesions and vice versa ^28^, while studies in humans have shown a correlation between Ab accumulation and basal forebrain atrophy in prodromal AD ^27,73^. Our findings suggest that the cholinergic-vascular hypothesis of disease could be extended to include disruption of glymphatic function not only in AD but also other disorders such as Parkinson’s disease dementia in which BFCN degeneration is a feature.

The strong correlation we observed between BFCN neuronal density and BOLD-CSF coupling and the correspondence between MS/VDB cholinergic ablation and reduced pulsation and glymphatic flux at the innervated artery demonstrate a region-specific regulation by BFCNs and their target brain area. As basal forebrain nuclei project to different cortical regions, whether differential rates of degeneration in these nuclei in prodromal AD ^74^ contribute to preferential deposition of pathological molecules in AD will need to be determined. In particular, Aβ deposition in sporadic AD starts from the default-mode network (DMN) that is important for cognitive functions ^75^. Previous studies have interpreted this association as Aβ impairment of functional connectivity. However, there may be a reverse association. Functional connectivity is regulated by BFCNs ^31^, with sustained improvement of DMN functional connectivity being reported in AD following AChEI treatment ^76^. Loss of BFCN function could therefore impair not only network activity but also glymphatic flux, leading to accelerated Aβ accumulation in this network. It will therefore be important to explore whether improving cerebrovascular function enhances BOLD-CSF coupling, arterial pulsation and glymphatic flux in humans to promote clearance of pathological molecules. In summary, our results highlight that BFCNs regulate glymphatic flux by modulating arterial pulsation, coordinating hemodynamic-CSF coupling, thereby leading to a change in fluid flux, and suggest that the cholinergic-vascular unit is a potential target for improving glymphatic function.

## Methods

### Human subjects

The human study was approved by The Prince Charles Hospital Human Research Ethics Committee. All participants were reviewed by a consultant geriatrician to determine those who met the Petersen criteria for MCI, including memory complaint, normal activities of daily living, normal general cognitive function with MMSE ≥ 24, abnormal memory for age and lack of dementia ^77^. Participants were excluded if there was evidence of neurological or psychiatric disorders (see our previous study ^39^ for detailed exclusion criteria). Informed written consent was obtained from participants. A blood sample, neuropsychological assessment and brain imaging were collected from each participant. A battery of cognitive assessments was conducted by an experienced neuropsychologist. Four cognitive domains were assessed: memory (including Rey Auditory Verbal Learning Test – short delay and long delay and Wechsler Memory Scale - Visual Reproduction I and II), executive function (including Trail-Making Test B, Controlled Oral Word Association Test, Wechsler Adult Intelligence Scale - Digit Span Backwards), attention (including Trail-Making Test A and Victoria Stroop Test) and language (including Boston Naming Test and Semantic Fluency Test).

### PET radiotracers

^18^F-florbetaben of 300 ± 10% MBq was used to measure Aβ deposition with a 20 min scan acquired starting at 90 min post-injection. ^18^F-FEOBV of 240 ± 10% MBq was used to measure cholinergic terminal integrity, with 30 min dynamic scans acquired at 180 min post-bolus injection. A static FEOBV Image was constructed by averaging the co-registered frames of the dynamic imaging data within a 20-min time window.

### PET-MRI acquisition

Brain imaging was conducted on a 3 Tesla PET-MRI scanner (Biograph mMR, Siemens Healthineers, Erlangen, Germany). Ultrashort echo-time MRI was conducted for attenuation correction. A T1-weighted 3D MPRAGE image was acquired with TR = 2.3 s, echo time (TE) = 2.26 ms, inversion time = 0.9 s, flip angle = 8°, 1 mm isotropic resolution, and matrix 256 x 240 x 192. A T2-weighted fluid attention inversion recovery (FLAIR) image was acquired with TR = 5 s, TE = 386 ms, flip angle = 120°, 1 mm isotropic resolution, and matrix 256 x 256 x 160. Resting-state fMRI was conducted by 2D gradient-echo echo-planar imaging (EPI) with TR = 2.68 s, TE = 30 ms, flip angle = 90°, 3 mm isotropic resolution, matrix size 72 x 72 x 42, and 446 repetitions.

### Human imaging data analysis

Data were processed using MATLAB (The MathWorks Inc.), FSL (v6, https://www.fmrib.ox.ac.uk/fsl), ANTs (http://picsl.upenn.edu/software/ants/) and AFNI (https://afni.nimh.nih.gov/). Due to technical issues, the FEOBV scan was not completed in 4 subjects and 2 did not complete the fMRI scan. After data quality control, another 2 subjects showed excessive motion in their fMRI scans. This resulted in 9 MCI and 10 control subjects with complete FEOBV and fMRI data. The PET data were analyzed based on the pipeline described in ^39^. The basal forebrain subregional volumes were estimated from the structural T1-weighted MRI scans based on the cytoarchitectonic map of the basal forebrain cholinergic nuclei ^78^. White matter hyperintensities were automatically quantified from the T2-weighted FLAIR images using the HyperIntensity Segmentation Tool ^79^.

For BOLD-CSF coupling, the fMRI data were motion corrected by AFNI 3dvolreg, and slice timing correction by AFNI 3dTshift, followed by second-order detrend, bandpass filtering to 0.01-0.1 Hz and 4 mm Gaussian smoothing using AFNI 3dTproject. Time points with frame-wise displacement larger than 0.35 mm were scrubbed to reduce motion artifacts. The first and the last 5 scans were discarded to avoid filter artifacts. As head motion tends to occur more frequent in later scans, only the initial 150 scans were used in the analysis. A CSF mask was manually defined in the most inferior slice to derive the CSF inflow signal. The cortical and sub-regional masks were defined based on the Automated Anatomical Labeling (AAL) atlas and were nonlinearly transformed from the template space to the subject space via structural T1-weighted MRI scans using ANTs. The averaged BOLD signals from these cortical masks were extracted to calculate the BOLD-CSF coupling using cross-correlation (MATLAB xcorr function, maximum time lag: ±10 scans) and Pearson’s correlation (MATLAB corr function).

### Animal study design

All animal experiments were approved by the Animal Ethics Committee of the University of Queensland. Two studies were conducted. In the first study, we determined the effects of cholinergic neurons on glymphatic and vascular functions by lesioning cholinergic neurons (n = 8 mice) in young C57BL/6 mice (female, weight=27±3g, age = 10-12 weeks) and compared to sham controls (n = 9). Two MRI sessions were conducted. The first to measure brain structure, arterial pulsation and resting-state fMRI, and the second to measure glymphatic function using a Gd contrast agent in a subset of mice (n = 5 from sham controls, and n = 6 from lesioned mice). The second study was to evaluate the effects of AChEI treatment on glymphatic function in 12 C57BL/6 mice (female, weight=21±1g, age = 10-12 weeks) with cholinergic lesions. Animals were housed under a 12 h–12 h light-dark cycle with ad libitum access to water and food. Where mice required separation for surgical recovery and welfare reasons, they were housed two per cage separated by a visual- and olfactory-permeable barrier.

### Surgery for cholinergic lesion

Mice were anesthetized with ketamine (100 mg/kg, i.p.) and the muscle relaxant xylazine (10 mg/kg, i.p.) and placed in a stereotaxic frame (David Kopf Instruments). After exposing the skull, a small bur hole was drilled at A-P, 1 mm; M-L, 0 mm from Bregma. Injections of murine-p75-saporin (mu-p75-SAP; 0.5 mg/ml; Advanced Targeting Systems) or control rabbit-IgG-saporin (Rb-IgG-SAP; 0.5 mg/ml) were performed using a calibrated glass micropipette through a Picospritzer® II (Parker Hannifin) into the border between the MS and VDB at D-V, −4.2 mm. In the first study, the toxin was infused at a rate of 0.4 μl/min (1.5 μl total volume), which resulted in a large amount of ablation. In the second study, the toxin concentration was reduced to 0.3 mg/ml to preserve more cholinergic neurons and was infused at a rate of 0.18 μl/min (1.0 μl total volume). After removing the needle, the hole was filled with bone wax and the skin sutured. The analgesic Torbugesic (2 mg/kg) and the antibiotic Baytril (5 mg/kg) were then injected subcutaneously with the latter being given until 3 days post-surgery. The MRI scans were conducted at least three weeks after lesion surgery.

### Surgery for cisterna magna cannulation

For the glymphatic MRI scan, mice were anesthetized with 2-3% isoflurane and secured in a stereotaxic frame. After making a small skin incision in the neck, the muscle layers were retracted and the cisterna magna was exposed. A mouse intrathecal catheter (32G, SAI), pre-filled with gadopentetic acid (Gd-DTPA) solution (Magnevist®, Bayer) was inserted at an angle of 45° relative to the mouse head into the center of the cisterna magna. The cyanoacrylate glue (3M) was dropped onto the dural membrane surrounding the cannula and then a mixture of dental cement (Vertex) was applied to fix the catheter in place. The mouse was then transported under anesthesia to the MRI scanner.

### Donepezil treatment

Donepezil solution (0.4 mg/ml) was freshly prepared on the day of the MRI experiment by dissolving donepezil hydrochloride (Sigma-Aldrich^®^, catalogue ID: D6821) in saline. Right after cisterna magna cannulation, a dosage of 2mg/kg donepezil was injected intraperitoneally before transporting to the MRI scanner.

### Animal MRI acquisition

Mice were maintained with 2-3% isoflurane in a mixture of O_2_ and air in a 1:2 ratio and an i.p. catheter was inserted. After the animal was secured in an MRI holder with ear and tooth bars, a bolus of 0.05mg/kg medetomidine (Troy Laboratories) was injected i.p. and maintained by a constant infusion of 0.1mg/kg/h medetomidine (i.p.) 10 min after the bolus, at which point the isoflurane was reduced to between 0.25-0.5%. The respiration rate and rectal temperature were continuously monitored (Model 1030, SAII), with the temperature maintained at 36.5-37^°^C by a water heater. The peripheral oxygen saturation (SpO_2_) and heart rate were monitored by a pulse oximeter (SAII) and respiration was monitored by a pressure sensor. The physiology was maintained within normal ranges, with SpO_2_ at 95-100%, heart rate at 150-350 beats per minute, and respiration rate at 80-110 breaths per minute throughout the MRI scans.

MRI was conducted on a 9.4T preclinical scanner (Biospec 9.4/30, Bruker BioSpin GmbH). Two sessions of scans were conducted in the first cohort of mice. In the first session, an 86mm volume transmitting coil with a 10 mm single-channel receiving surface coil (Bruker) was used to acquire the structural MRI, fMRI, angiography and arterial pulsation imaging. In the second session, a 20 mm receiving surface coil (Bruker) was used for the glymphatic MRI and angiography for better signal uniformity across the brain. For the second cohort of mice, the above scans were conducted in one session using a 10 mm surface receive coil.

Localized high-order shim (MAPshim) based on the B_0_ map was applied. Structural T2-weighted MRI scans of 0.1×0.1×0.3 mm^3^ resolution, field of view (FOV) = 19.2 x 19.2 x 16.8 mm^3^ was acquired using a 2D fast spin-echo with TR/TE = 5500/40 ms and five averages. Resting-state fMRI was conducted using a multiband gradient-echo EPI with TR = 300 ms, TE = 15 ms, 16 slices with 4 bands, resolution = 0.3 x 0.3 x 0.6 mm^3^, and 2000 repetitions ^41^. Arterial pulsation was measured by a single-shot gradient-echo EPI of 2 horizontal slices with thickness = 0.5 mm, 2.5 mm gap, in-plane resolution = 0.2 x 0.2 mm^2^, FOV = 19.2 x 12.8 mm, matrix size = 96 x 64, TR/TE = 70/14.15 ms, and flip angle = 90° in order to capture the heart rate. A total of 2500 time frames were acquired in 2 min 55s. Following the surgery for cisterna magna cannulation, glymphatic imaging was acquired using dynamic 3D T1-weighted fast low angle shot (FLASH) MRI (TR/TE = 21/2.66ms, flip angle = 20°, matrix = 192 x 128 x 80, 0.1 mm isotropic resolution, and 3.33 min per volume) covering the whole brain. After 10 min of baseline scanning, 8.0 μl Gd-DTPA at 0.56 mmol/kg dosage was slowly infused into the cisterna magna at a rate of 0.5 μl/min for 16 min using an infusion pump. 40 volumes were continuously acquired for the total of 132 min in the first study and 50 volumes were acquired in the second study. Angiography was acquired using a time-of-flight (ToF) sequence with TR/TE = 17/3 ms, flip angle = 20°, slice number = 80, thickness = 0.35 mm, FOV = 20×20 mm^2^, matrix size = 320×320, and in-plane resolution = 0.0625 x 0.0625 mm^2^.

### Animal MRI data processing

Due to scanner or image quality issues, fMRI from 2 sham control mice and pulsation MRI from 4 lesioned mice were discarded. Due to blockage of the catheter for Gd injection, 1 control mouse in the first study and 1 vehicle-treated and 1 donepezil-treated mouse in the second study did not achieve Gd contrast and were removed from the glymphatic analysis.

For BOLD-CSF coupling analysis, the same data processing steps as those used in human fMRI were applied, with additional EPI distortion correction by FSL Topup and baseline intensity normalization of the voxel time-series to convert the fMRI signal to percentage signal change. A Gaussian smoothing of 0.6 mm was applied and 250 scans were discarded from each end to avoid filter artifacts. Skull stripping was performed automatically by PCNN3D ^80^ (https://sites.google.com/site/chuanglab/software/3d-pcnn) and then manually inspected and edited. Cortical and hippocampal masks were derived from the Australian Mouse Brain Mapping Consortium (AMBMC; http://www.imaging.org.au/AMBMC/AMBMC) atlas and transformed to the subject space via the T2-weighted structural MRI. Manual editing was conducted to avoid overlapping with the ventricles. The CSF mask was defined in the posterior slices near the cerebellum.

For glymphatic MRI, the Gd-enhanced time-series image was denoised by MP-PCA ^81^, motion corrected and normalized by the mean intensity of the 3 baseline scans in each voxel. The AUC of each voxel, calculated by summing the intensity after the Gd injection, was used to represent the total glymphatic flow volume. The AUC signal of the whole brain, which represents the effective amount of contrast agent delivered into the brain, was used to normalize the signal change in each voxel to account for individual variation. To estimate the glymphatic flux, the Gd contrast arrival time was defined as the time when the signal increased above 20% of the maximum signal change. The signal rising slope between the arrival time and time-to-peak was used to represent the influx rate. The signal decay time was derived from the signal change after the peak signal by a 3-parameter fitting to an exponential function: *Ae^−t/Decay^ + C* (MATLAB fit function). For voxel-wise group comparison, linear and non-linear transformations were applied with ANTs, to register the data to the AMBMC brain template. Study-specific templates were made from the registered images from all animals. A second round of linear and non-linear transformation was then estimated to register the data to the study-specific template. The transformation was then applied to the AUC and kinetic parameter maps using linear interpolation. A 0.2-mm Gaussian smoothing was applied to the co-registered maps to account for residual misregistration.

In the pulsation MRI scans, the time-course signal of each pixel was first normalized by its temporal mean and then Fourier transformed. The temporal standard deviation of the normalized time-course was calculated to map the amplitude of signal fluctuation, which revealed that the highest fluctuation colocalized with cerebral arteries. We manually drew 5 region-of-interests (ROIs), two for the PCA, one for the anterior cerebral artery (ACA) and 2 for the LHiA. Based on the spectra and the heart rates recorded, a bandpass filter of 2 Hz width around the cardiac peak, which encompassed the heart rate and the aliased first harmonics, was applied. The temporal variance of the filtered signal was used as a measure of arterial pulsation power.

### Tissue processing and immunohistochemistry

Immediately after the glymphatic MRI, animals were deeply anesthetized with sodium pentobarbitone and transcardially perfused with ice-cold phosphate-buffered saline (PBS), followed by 4% paraformaldehyde in PBS (pH 7.4). Brains were removed, post-fixed overnight at 4°C in 4% paraformaldehyde and then placed in 30% sucrose solution. Serial brain coronal sections were cut at a thickness of 40 μm and placed in a six-series configuration. All sections were washed in PBS and stored at 4°C in PBS + 0.01% sodium azide until used.

Sections were washed three times with PBS prior to incubation with blocking solution (0.1M PBS/0.1% Triton-×100 /5% normal horse serum/0.05% sodium azide) for 80 min at room temperature. Basal forebrain tissue sections of each brain were selected based on the anatomical features of corpus callosum and anterior commissure. Immunofluorescence labeling of the cholinergic cells were stained with primary goat anti-p75 antibody (1:400, AF1157, R&D Systems) overnight at room temperature. After washing these slices in 0.1% Triton-X in 0.1M PBS, the sections were incubated at room temperature with anti-goat IgG Alexa Fluor 488 secondary antibody (1:1000, A11055, Thermo Fisher Scientific) and 4’, 6-diamidino-2-phenylindole, dihydrochloride (DAPI) (1:5000, D9542, Sigma-Aldrich). After washing, sections were then mounted onto slides using fluorescence mounting medium (Dako).

### Histological analysis and quantification

Images of histological sections were obtained using an upright fluorescence slide scanner (Zeiss Axio Imager Z2) with a 20x objective and AxioVision software (Carl Zeiss). The basal forebrain cell counts were performed in slices from 1.3 to 0.1mm anterior to Bregma, with every third section (a total of 10 sections per animal over 120μm) being analyzed. All measurements and analyses were performed using Imaris (software ver 7.2.3, Bitplane Co.). Positive p75 and DAPI staining within the basal forebrain was quantified section-by-section by the ‘spots’ function in Imaris. The borders of the MS and VDB were manually drawn in accordance with the Paxinos and Franklin mouse brain atlas (fifth edition), and the numbers of p75- and DAPI-positive cells were quantified within each ROI, then were normalized to the area of the ROI to obtain the cholinergic neuron and general cell density.

### Statistical analysis

Voxel-wise comparison of MRI data was conducted using 2-sample t-tests (AFNI 3dttest++), with a voxel level threshold at p<0.05 and cluster-level correction of multiple comparison (p<0.05) by AFNI 3DClustSim. A voxel-wise correlation with the cholinergic neuron count was calculated by AFNI 3dTcorr1D, and thresholded by the same method as above. Statistical analysis of regional data was performed using the MATLAB Statistics Toolbox. Between-group differences were compared using two-sample t-tests or non-parametric Wisconsin rank-sum tests, with significance set at p < 0.05. Linear regression was calculated using MATLAB fitlm function. A Pearson’s or Spearman’s correlation analysis was conducted between neuropsychological data, cell densities and imaging measurements, with p < 0.05 being considered significant. Values are expressed as the mean ± standard error of the mean (SEM).

## Data availability

Source data for quantifications shown in all graphs plotted in figures and Extended Data figures are available in the online version of the paper. The raw data acquired in this study are also available from the corresponding authors upon reasonable request.

## Code availability

Custom code used in the current study is available from the corresponding authors upon reasonable request.

## Acknowledgements

We thank Rowan Tweedale for proof reading and Dr Yunpeng Wang for setting up the glymphatic surgery and imaging. The study was supported by the National Health and Medical Research Council (NHMRC), Grantor Reference ID: GNT1162505 to K.H.C and E.J.C; the Dementia Australia Research Foundation Project Grant ID: RM2022001731 to Z.L.; and the Australian Research Council (ARC) Discovery Project grant ID: DP240101321 to K.H.C. The human study was funded by The Common Good Foundation, an initiative of The Prince Charles Hospital Foundation 2014 Program Grant: PRO2014-10, and Bupa Foundation to E.E. and E.C.

## Author contributions

K.H.C. designed the study, designed and supervised MRI experiments, analyzed and interpreted the data, wrote the manuscript, and directed the project. X.A.Z., L.Q. and Z.L. performed the animal experiments and wrote the manuscript; G.N. and Z.L. performed the histology and analyzed the data; Y.X. E.E. and J.F. designed the human experiments and processed the human data; E.J.C. designed the study, interpreted the data, and wrote the manuscript.

## References

1. Nedergaard, M. et al. Glymphatic failure as a final common pathway to dementia. Science (80-.). 56, 50–56 (2020).

2. Ahn, J. H. et al. Meningeal lymphatic vessels at the skull base drain cerebrospinal fluid. Nature 572, 62–66 (2019).

3. Arbel-Ornath, M. et al. Interstitial fluid drainage is impaired in ischemic stroke and Alzheimer’s disease mouse models. Acta Neuropathol. 126, 353–364 (2013).

4. Iliff, J. J. et al. A paravascular pathway facilitates CSF flow through the brain parenchyma and the clearance of interstitial solutes, including amyloid β. Sci. Transl. Med. 4, 147ra111 (2012).

5. Iliff, J. J. et al. Cerebral arterial pulsation drives paravascular CSF-interstitial fluid exchange in the murine brain. J. Neurosci. 33, 18190–9 (2013).

6. Mestre, H. et al. Flow of cerebrospinal fluid is driven by arterial pulsations and is reduced in hypertension. Nat. Commun. 9, 4878 (2018).

7. Fultz, N. E. et al. Coupled electrophysiological, hemodynamic, and cerebrospinal fluid oscillations in human sleep. Science (80-.). 366, 628–631 (2019).

8. Jiang-Xie, L.-F. et al. Neuronal dynamics direct cerebrospinal fluid perfusion and brain clearance. Nature 627, 157–164 (2024).

9. Kress, B. T. et al. Impairment of paravascular clearance pathways in the aging brain. Ann. Neurol. 76, 845–861 (2014).

10. Eide, P. K. et al. Sleep deprivation impairs molecular clearance from the human brain. Brain 144, 863–874 (2021).

11. Da Mesquita, S. et al. Functional aspects of meningeal lymphatics in ageing and Alzheimer’s disease. Nature 560, 185–191 (2018).

12. Holth, J. K. et al. The sleep-wake cycle regulates brain interstitial fluid tau in mice and CSF tau in humans. Science (80-.). 363, 880–884 (2019).

13. Harrison, I. F. et al. Impaired glymphatic function and clearance of tau in an Alzheimer’s disease model. Brain 143, 2576–2593 (2020).

14. Ding, X.-B. et al. Impaired meningeal lymphatic drainage in patients with idiopathic Parkinson’s disease. Nat. Med. 27, 411–418 (2021).

15. Rasmussen, M. K. et al. The glymphatic pathway in neurological disorders. Lancet Neurol. 17, 1016–1024 (2018).

16. van Veluw, S. J. et al. Vasomotion as a driving force for paravascular clearance in the awake mouse brain. Neuron 105, 549–561.e5 (2020).

17. Hablitz, L. M. et al. Increased glymphatic influx is correlated with high EEG delta power and low heart rate in mice under anesthesia. Sci. Adv. 5, eaav5447 (2019).

18. Xie, L. et al. Sleep drives metabolite clearance from the adult brain. Science 342, 373– 7 (2013).

19. Holstein-Rønsbo, S. et al. Glymphatic influx and clearance are accelerated by neurovascular coupling. Nat. Neurosci. 26, 1042–1053 (2023).

20. Gouveia-Freitas, K. et al. Perivascular spaces and brain waste clearance systems: relevance for neurodegenerative and cerebrovascular pathology. Neuroradiology 63, 1581–1597 (2021).

21. Drieu, A. et al. Parenchymal border macrophages regulate the flow dynamics of the cerebrospinal fluid. Nature 611, 585–593 (2022).

22. Mestre, H. et al. Periarteriolar spaces modulate cerebrospinal fluid transport into brain and demonstrate altered morphology in aging and Alzheimer’s disease. Nat. Commun. 13, 3897 (2022).

23. Hampel, H. et al. The cholinergic system in the pathophysiology and treatment of Alzheimer’s disease. Brain 141, 1917–1933 (2018).

24. Hamel, E. Perivascular nerves and the regulation of cerebrovascular tone. J. Appl. Physiol. 100, 1059–1064 (2006).

25. Hamel, E. Cholinergic modulation of the cortical microvascular bed. Prog. Brain Res. 145, 171–178 (2004).

26. Potter, P. E. et al. Pre- and post-synaptic cortical cholinergic deficits are proportional to amyloid plaque presence and density at preclinical stages of Alzheimer’s disease. Acta Neuropathol. 122, 49–60 (2011).

27. Kerbler, G. M. et al. Basal forebrain atrophy correlates with amyloid β burden in Alzheimer’s disease. NeuroImage. Clin. 7, 105–13 (2015).

28. Ramos-Rodriguez, J. J. et al. Rapid β-amyloid deposition and cognitive impairment after cholinergic denervation in APP/PS1 mice. J. Neuropathol. Exp. Neurol. 72, 272– 285 (2013).

29. Nizari, S. et al. Loss of cholinergic innervation differentially affects eNOS-mediated blood flow, drainage of Aβ and cerebral amyloid angiopathy in the cortex and hippocampus of adult mice. Acta Neuropathol. Commun. 9, 12 (2021).

30. German-Castelan, L. et al. Sex-dependent cholinergic effects on amyloid pathology: A translational study. Alzheimer’s Dement. 20, 995–1012 (2024).

31. Turchi, J. et al. The basal forebrain regulates global resting-state fMRI fluctuations. Neuron 97, 940–952.e4 (2018).

32. Liu, X. et al. Subcortical evidence for a contribution of arousal to fMRI studies of brain activity. Nat. Commun. 9, 395 (2018).

33. Petrou, M. et al. In Vivo Imaging of Human Cholinergic Nerve Terminals with (-)-5-18F-Fluoroethoxybenzovesamicol: Biodistribution, Dosimetry, and Tracer Kinetic Analyses. J. Nucl. Med. 55, 396–404 (2014).

34. Han, F. et al. Reduced coupling between cerebrospinal fluid flow and global brain activity is linked to Alzheimer disease-related pathology. PLoS Biol. 19, e3001233 (2021).

35. Han, F. et al. Resting-state global brain activity affects early β-amyloid accumulation in default mode network. Nat. Commun. 14, 7788 (2023).

36. Zhang, Y. et al. Reduced coupling between the global blood-oxygen-level-dependent signal and cerebrospinal fluid inflow is associated with the severity of small vessel disease. NeuroImage Clin. 36, 103229 (2022).

37. Ferdinando, H. et al. Altered cerebrovascular-CSF coupling in Alzheimer’s disease measured by functional near-infrared spectroscopy. Sci. Rep. 13, 22364 (2023).

38. Klunk, W. E. et al. The Centiloid Project: standardizing quantitative amyloid plaque estimation by PET. Alzheimers. Dement. 11, 1–15.e1–4 (2015).

39. Xia, Y. et al. Reduced cortical cholinergic innervation measured using [18F]-FEOBV PET imaging correlates with cognitive decline in mild cognitive impairment. NeuroImage. Clin. 34, 102992 (2022).

40. Zaborszky, L. et al. The basal forebrain cholinergic projection system in mice. in The Mouse Nervous System (eds. Watson, C., Paxinos, G. & Puelles, L.) 684–718 (Elsevier Inc., 2012). doi:10.1016/B978-0-12-369497-3.10028-7.

41. Lee, H.-L. et al. Ultrafast fMRI of the rodent brain using simultaneous multi-slice EPI. Neuroimage 195, 48–58 (2019).

42. Iliff, J. J. et al. Brain-wide pathway for waste clearance captured by contrast-enhanced MRI. J. Clin. Invest. 123, 1299–1309 (2013).

43. Benveniste, H. et al. Glymphatic cerebrospinal fluid and solute transport quantified by MRI and PET imaging. Neuroscience 474, 63–79 (2021).

44. Benveniste, H. et al. Anesthesia with dexmedetomidine and low-dose isoflurane increases solute transport via the glymphatic pathway in rat brain when compared with high-dose isoflurane. Anesthesiology 127, 976–988 (2017).

45. Ballinger, E. C. et al. Basal forebrain cholinergic circuits and signaling in cognition and cognitive decline. Neuron 91, 1199–1218 (2016).

46. Xu, M. et al. Basal forebrain circuit for sleep-wake control. Nat. Neurosci. 18, 1641– 1647 (2015).

47. Nizari, S. et al. 3D reconstruction of the neurovascular unit reveals differential loss of cholinergic innervation in the cortex and hippocampus of the adult mouse brain. Front. Aging Neurosci. 11, 172 (2019).

48. Xiong, B. et al. Precise cerebral vascular atlas in stereotaxic coordinates of whole mouse brain. Front. Neuroanat. 11, 1–17 (2017).

49. Gao, J. H. et al. Quantitative assessment of blood inflow effects in functional MRI signals. Magn. Reson. Med. 36, 314–319 (1996).

50. Mirsky, I. Pulse velocities in cylindrical, tapered and curved anisotropic elastic arteries. Bull. Math. Biol. 35, 495–511 (1973).

51. Webb, A. J. S. et al. Magnetic resonance imaging measurement of transmission of arterial pulsation to the brain on propranolol versus amlodipine. Stroke 47, 1669–1672 (2016).

52. Hamlin, A. S. et al. Lesions of the basal forebrain cholinergic system in mice disrupt idiothetic navigation. PLoS One 8, e53472 (2013).

53. Holschneider, D. P. et al. Changes in electrocortical power and coherence in response to the selective cholinergic immunotoxin 192 IgG-saporin. Exp. Brain Res. 126, 270– 280 (1999).

54. Fok, H. et al. Regulation of vascular tone and pulse wave velocity in human muscular conduit arteries. Hypertension 60, 1220–1225 (2012).

55. Peng, W. et al. Suppression of glymphatic fluid transport in a mouse model of Alzheimer’s disease. Neurobiol. Dis. 93, 215–225 (2016).

56. Taoka, T. et al. Evaluation of glymphatic system activity with the diffusion MR technique: diffusion tensor image analysis along the perivascular space (DTI-ALPS) in Alzheimer’s disease cases. Jpn. J. Radiol. 35, 172–178 (2017).

57. Vijayakrishnan Nair, V., et al. Neurofluid coupling during sleep and wake states. Sleep Med. 110, 44–53 (2023).

58. Wang, Y. et al. Cerebrovascular activity is a major factor in the cerebrospinal fluid flow dynamics. Neuroimage 258, 119362 (2022).

59. Lee, M. G. et al. Hippocampal theta activity following selective lesion of the septal cholinergic systeM. Neuroscience 62, 1033–1047 (1994).

60. Lecrux, C. et al. Impact of altered cholinergic tones on the neurovascular coupling response to whisker stimulation. J. Neurosci. 37, 1518–1531 (2017).

61. Chaudhary, S. et al. Hemodynamic effects of cholinesterase inhibition in mild Alzheimer’s disease. J. Magn. Reson. Imaging 38, 26–35 (2013).

62. Nakano, S. et al. Donepezil hydrochloride preserves regional cerebral blood flow in patients with Alzheimer’s disease. J. Nucl. Med. 42, 1441–1445 (2001).

63. Li, W. et al. Changes in regional cerebral blood flow and functional connectivity in the cholinergic pathway associated with cognitive performance in subjects with mild Alzheimer’s disease after 12-week donepezil treatment. Neuroimage 60, 1083–1091 (2012).

64. Pa, J. et al. Cholinergic enhancement of functional networks in older adults with mild cognitive impairment. Ann. Neurol. 73, 762–73 (2013).

65. Han, G. et al. Arterial pulsation dependence of perivascular cerebrospinal fluid flow measured by dynamic diffusion tensor imaging in the human brain. Neuroimage 297, 120653 (2024).

66. Wen, Q. et al. Assessing pulsatile waveforms of paravascular cerebrospinal fluid dynamics using dynamic diffusion-weighted imaging (dDWI). Neuroimage 260, 119464 (2022).

67. Driver, I. D. et al. Most Small Cerebral Cortical Veins Demonstrate Significant Flow Pulsatility: A Human Phase Contrast MRI Study at 7T. Front. Neurosci. 14, 415 (2020).

68. Hermes, D. et al. Measuring brain beats: Cardiac-aligned fast functional magnetic resonance imaging signals. Hum. Brain Mapp. 44, 280–294 (2023).

69. Shearman, E. et al. Changes in cerebral neurotransmitters and metabolites induced by acute donepezil and memantine administrations: A microdialysis study. Brain Res. Bull. 69, 204–213 (2006).

70. Claassen, J. A. H. R. et al. Cholinergically mediated augmentation of cerebral perfusion in Alzheimer’s disease and related cognitive disorders: The cholinergic-vascular hypothesis. Journals Gerontol. - Ser. A Biol. Sci. Med. Sci. 61, 267–271 (2006).

71. Siow, T. Y. et al. Association of sleep, neuropsychological performance, and gray matter volume with glymphatic function in community-dwelling older adults. Neurology 98, e829–e838 (2022).

72. Schmitz, T. W. et al. Basal forebrain degeneration precedes and predicts the cortical spread of Alzheimer’s pathology. Nat. Commun. 7, 13249 (2016).

73. Teipel, S. et al. Cholinergic basal forebrain atrophy predicts amyloid burden in Alzheimer’s disease. Neurobiol. Aging 35, 482–491 (2014).

74. Kilimann, I. et al. Subregional basal forebrain atrophy in alzheimer’s disease: A multicenter study. J. Alzheimer’s Dis. 40, 687–700 (2014).

75. Palmqvist, S. et al. Earliest accumulation of β-amyloid occurs within the default-mode network and concurrently affects brain connectivity. Nat. Commun. 8, 1214 (2017).

76. Blautzik, J. et al. Functional connectivity increase in the default-mode network of patients with Alzheimer’s disease after long-term treatment with galantamine. Eur. Neuropsychopharmacol. 26, 602–613 (2016).

77. Petersen, R. C. et al. Mild cognitive impairment: Clinical characterization and outcome. Arch. Neurol. 56, 303–308 (1999).

78. Zaborszky, L. et al. Stereotaxic probabilistic maps of the magnocellular cell groups in human basal forebrain. Neuroimage 42, 1127–1141 (2008).

79. Manjón, J. V. et al. MRI white matter lesion segmentation using an ensemble of neural networks and overcomplete patch-based voting. Comput. Med. Imaging Graph. 69, 43–51 (2018).

80. Chou, N. et al. Robust automatic rodent brain extraction using 3-D pulse-coupled neural networks (PCNN). IEEE Trans. Image Process. 20, 2554–64 (2011).

81. Veraart, J. et al. Denoising of diffusion MRI using random matrix theory. Neuroimage 142, 394–406 (2016).

